# Across-subject offline decoding of motor imagery from MEG and EEG

**DOI:** 10.1101/349225

**Authors:** Hanna-Leena Halme, Lauri Parkkonen

**Affiliations:** Department of Neuroscience and Biomedical Engineering NBE, Aalto University School of Science, P.O. Box 12200, FI-00076 Aalto, Finland; Aalto Neuroimaging, MEG Core, Aalto University School of Science, Espoo, Finland

**Author notes:** Corresponding author: Hanna-Leena Halme Department of Neuroscience and Biomedical Engineering NBE, Aalto University School of Science, P.O. Box 12200, FI-00076 Aalto, Finland.

## Abstract

Long calibration time hinders the feasibility of brain-computer interfaces (BCI). If other subjects’ data were used for training the classifier, BCI-based neurofeedback practice could start without the initial calibration. Here, we compare methods for inter-subject decoding of left- vs. right-hand motor imagery (MI) from MEG and EEG.

Six methods were tested on data involving MEG and EEG measurements of healthy participants. Only subjects with good within-subject accuracies were selected for inter-subject decoding. Three methods were based on the Common Spatial Patterns (CSP) algorithm, and three others on logistic regression with l_1_ - or l_2,1_ -norm regularization. The decoding accuracy was evaluated using 1) MI and 2) passive movements (PM) for training, separately for MEG and EEG.

When the classifier was trained by MI, the best accuracies across subjects (mean 70.6% for MEG, 67.7% for EEG) were obtained using multi-task learning (MTL) with logistic regression and l_2_,_1_-norm regularization. MEG yielded slightly better average accuracies than EEG. When PM were used for training, none of the inter-subject methods yielded above chance level (58.7%) accuracy.

In conclusion, MTL and training with other subject’s MI is efficient for inter-subject decoding of MI. Passive movements of other subjects are likely suboptimal for training the MI classifiers.

## 1. INTRODUCTION

Brain–computer interfaces (BCI) translate user’s brain activity in real time into commands for an external device. Non-invasive BCIs have diverse applications in both basic and clinical neuroscience, including videogames, exercise systems for increasing attention or relaxation, communication devices for disabled people and various tools for neurological rehabilitation. For example, hemiparetic patients can learn to control an orthosis attached to the paretic hand using motor imagery (MI) and concurrent feedback (see a review by Teo and co-workers ^1^). This so-called neurofeedback training may enhance recovery of sensorimotor function compared to traditional physiotherapy, but the clinical evidence of BCI therapy still remains vague ^2,3^.

Electric brain activity can be measured noninvasively with e.g. electroencephalography (EEG) and the corresponding magnetic fields with magnetoencephalography (MEG). MEG has recently gained interest in the context of BCI experiments as well ^4–7^, mainly because it provides higher signal-to-noise ratio and spatial resolution compared to EEG. MEG can be used to assist the development of eventually EEG-based BCIs, and MEG-based BCIs can be used in basic neuroscientific research.

Unfortunately, BCIs typically require dozens or even hundreds of samples of subject’s neurophysiological signals for training the classification algorithms to decode the user’s brain states with an accuracy exceeding the chance level. Due to large inter-subject variability of these signals, the user-specific brain responses are typically collected in the beginning of each session for calibrating the BCI and thus the total calibration time can be significant. This process can be exhausting for patients suffering from neurological disorders. Although Foldes and co-workers showed previously that five minutes of MEG data are sufficient for calibrating a subject-specific BCI discriminating MI from rest ^7^, this time is still away from therapy. In addition, when employing neurofeedback for rehabilitation, the feedback should drive brain activity towards that of a healthy brain and not to reinforce the prevailing pathological state as a patient-specific BCI might do.

In order to overcome these drawbacks, several research groups have made effort to develop intersubject-generalized BCIs ^8–10^ which can be trained in advance using data from previously measured subjects. Therefore, a new BCI user can begin the neurofeedback practice immediately without the initial calibration session, which saves both time and effort of the patients.

Successful inter-subject classification requires that globally relevant signal features are extracted from each training subject. Optimal features should accurately discriminate between the brain states of interest and they should generalize across subjects in order to enable inter-subject learning. Feature extraction is a crucial part of a BCI, and selecting optimal features can significantly improve the classification accuracy, as shown e.g. in our previous study ^11^.

Multi-task learning (MTL ^12,13^) aims at selecting a few relevant features from a large feature set such that these features are shared across multiple related tasks; in the case of BCI training, the tasks are usually measurements from different subjects. MTL is beneficial for learning problems in which the number of samples is much lower than the number of features, since it effectively reduces the dimensionality of the feature space. MTL finds a sparse feature set that is correlated across tasks instead of optimizing features for each individual task, which makes the method robust to outliers. In case of inter-subject classification, MTL considers decoding of each subject’s data as a separate task. In contrast to data pooling, MTL does not assume that training data from all subjects are equally distributed, which allows variability between subjects.

Regularized logistic regression has been used, among other machine-learning applications, for classifying EEG signals^14,15^. Moreover, logistic regression with l_2,1_- norm regularization is found to be efficient for selecting a small number of relevant features from a high-dimensional feature space ^16^. Recently, MTL based on logistic regression and regularization with l_2,1_ norm minimization was successfully used for extracting features and classifying MEG signals in an inter-subject decoding paradigm ^17^.

Another popular method for feature extraction is spatial filtering with the Common Spatial Patterns (CSP) algorithm, which has been applied to EEG ^18,19^ as well as to MEG ^11^. CSP aims at maximizing the covariance of signals representing one class (e.g. left-hand motor imagery) while simultaneously minimizing it for another class (e.g. right-hand motor imagery). The benefits of CSP are low computational cost and efficiency especially in motor function decoding^20^, but the extension to inter-subject classification is challenging, since the spatial patterns of each individual are slightly different, which may result in classifier overfitting.

In order to avoid overfitting and to improve generalization, both the covariance matrices and the objective function of CSP can be regularized towards a more generic filter decomposition by using additional regularization parameters ^21^. Regularization of CSP has been found to improve inter-subject classification of EEG data ^8,21–23^, but to our knowledge has not yet been applied to MEG data. Furthermore, the discriminatory power of CSP can be improved with preceding dimensionality reduction using e.g. Spatio-Spectral Decomposition (SSD^24^), which maximizes the signal power at a frequency of interest while simultaneously minimizing it at the flanking frequencies. SSD is especially beneficial for extracting neural oscillations occurring within a known frequency band, such as sensorimotor rhythms (SMR) consisting of ~10-Hz and ~20-Hz oscillations.

In the current study, we investigate inter-subject decoding of motor imagery (MI) -related MEG and EEG signals using different methods for feature selection and classification. The classification accuracy is evaluated using a dataset from healthy participants. The data include both MEG and EEG collected during passive movements (PM) and MI of the left and right hand. The decoding accuracy is evaluated using 1) MI and 2) PM as training data, separately for MEG and EEG. The rationale for using other subjects’ PM for training is based on previous studies showing that PM of the test subject can be used for calibration of an MI-BCI^25–27^. However, there is also evidence that the activity patterns during MI and PM are not similar^28^, which likely reduces the accuracy in inter-subject MI classification. Moreover, in previous studies, the decoding was performed between rest and MI, whereas in the current study we are decoding left- vs. right-hand MI. We therefore expect inferior results with PM training compared to MI training.

The evaluated decoding methods are based on either 1) CSP combined with linear discriminant analysis (LDA) classifier, or 2) logistic regression with l_2,1_- or l_1_-norm regularization. For the CSP-based methods, the following approaches (abbreviation in parentheses) are used for decoding:

a. CSP, classification with LDA (CSP+LDA)
b. CSP, classification with bootstrap aggregating & LDA (CSP+bagging)
c. Regularized CSP, classification with LDA (regCSP)

The functions included in MTJFL^17^ and MALSAR ^29^ Matlab toolboxes were modified and used for the logistic regression –based methods. The following inter-subject decoding approaches, implemented in the MTJFL toolbox, are used for decoding:

a. Pooling with logistic regression and l_1_-norm regularization (pooling)
b. MTL with logistic regression and l_1_-norm regularization (L1-MTL)
c. MTL with logistic regression and l_2,1_-norm regularization (L21-MTL)

In addition, we calculate the accuracies for within-subject decoding for comparison with the inter-subject decoding results. In this case, each subject’s own PM or MI data are used for training. PM training for MEG is tested in an online BCI paradigm using CSP+LDA. For EEG, similar PM training paradigm is conducted for the offline data. For both modalities, within-subject MI training is calculated offline using 10-fold cross-validation and l_1_-norm regularized logistic regression.

The main research questions of this study are: 1) do generalized classifiers perform above chance level and how does their performance compare to that of a subject-specific classifier, 2) can generalized classifiers be trained equally well with responses to PM and MI, and 3) whether MEG and EEG data yield different inter-subject decoding results.

## 2. RESULTS

The detailed results for different modalities and training data are presented in the following subsections. All classification results are summarized in Fig. 1.

**Figure 1.**
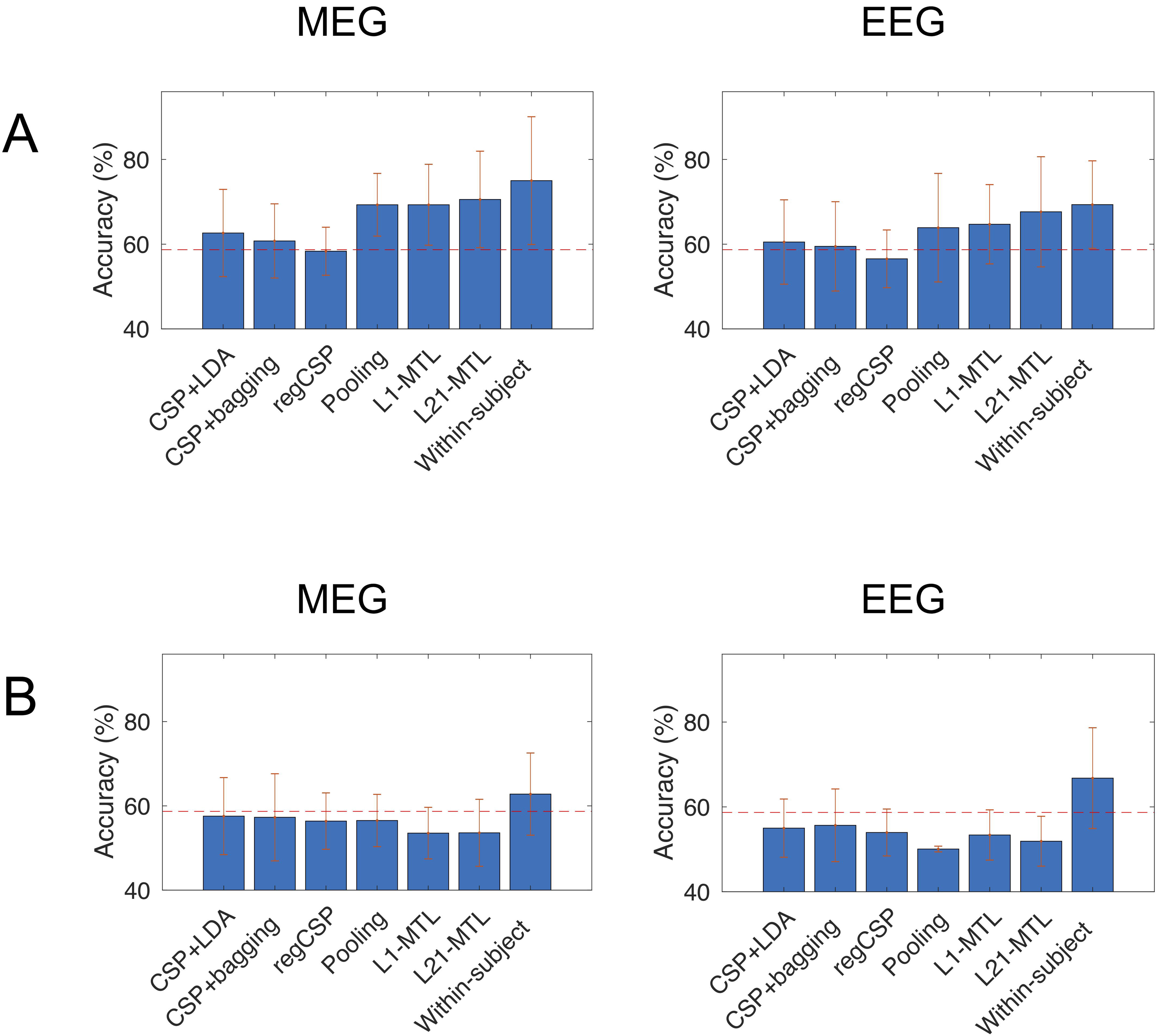
Classification results for (A) Motor-imagery-trained MEG and EEG, (B) Passive-movement-trained MEG and EEG. The chance level is indicated with a dashed line. Error bars represent the standard deviation.

### 2.1. Within-subject decoding online and offline

Table 1 represents the results for within-subject MEG decoding 1) in the online BCI paradigm, using subject’s own PM for training (see Section 4.4 for a detailed explanation) and 2) in the offline analysis using 10-fold cross-validation with subject’s own MI data and l_1_-norm regularized logistic regression.

**Table 1.**
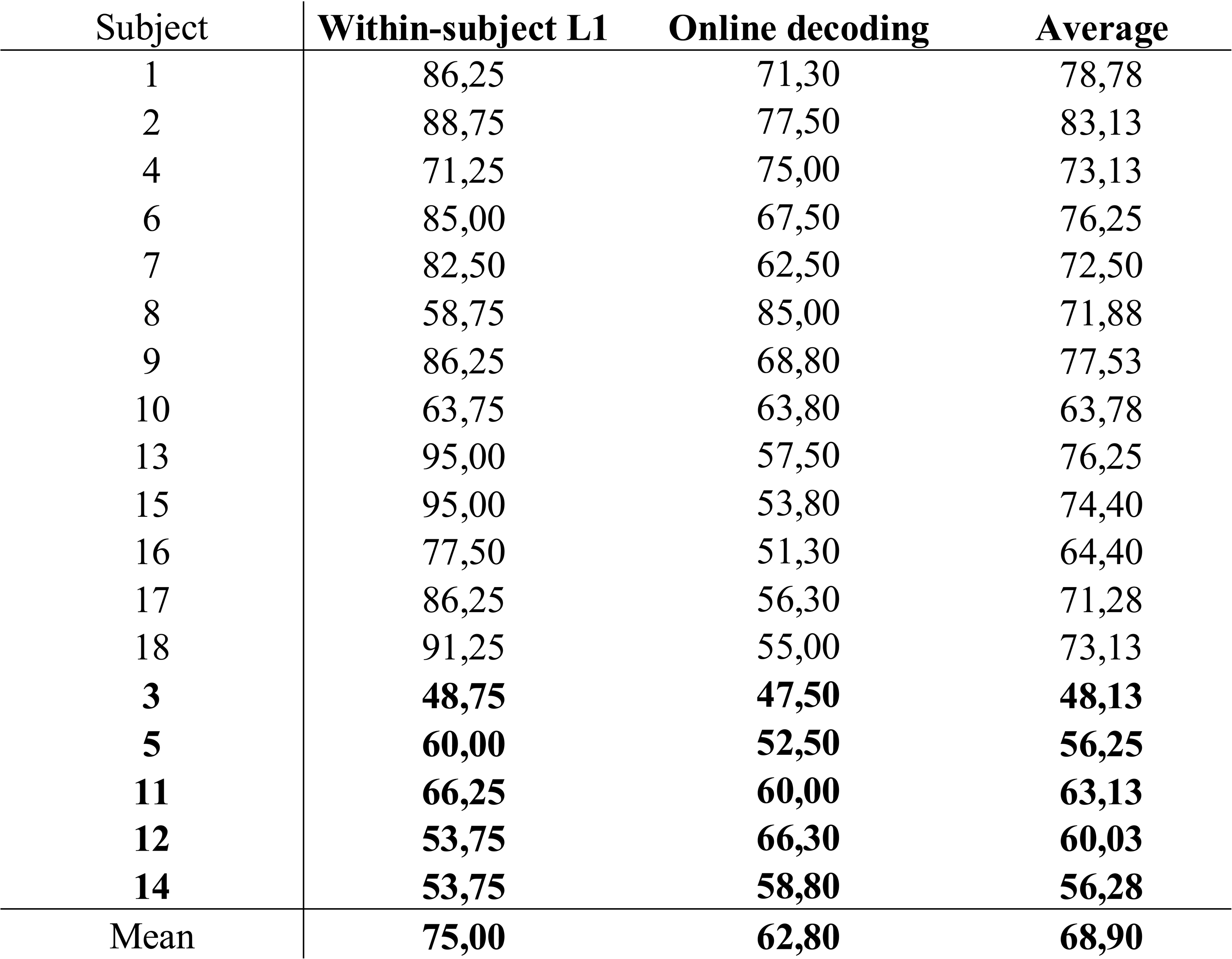
Accuracies for offline 10-fold cross-validation with MI training and online PM-trained decoding for individual subjects. The subjects discarded from the training data for inter-subject classification are written in bold.

The online accuracy was below the sample-size-adjusted chance level of 58.7%^30^ for 7 subjects. However, as the online decoding was done with PM training, we assumed these results probably did not reflect the robustness of MI or the ability to perform MI. Therefore, the decoding accuracy was also calculated offline with MI training in order to evaluate the individual MI performance. The mean accuracy for each subject was calculated as the average of online and offline within-subject accuracies. Five subjects with the lowest mean accuracy were discarded from the training data in further analyses, i.e. the poorly performing subjects were used only for testing the decoders. The purpose of discarding bad performers from the training data was to find out whether their results would be improved with good-quality training data from good performers.

### 2.2. Offline training by motor imagery

Table 2 shows the results when the classifier was trained by MEG responses to motor imagery; correponding results for EEG are shown in Table 3. The best inter-subject mean accuracies of 70.6% for MEG and 67.7% for EEG were obtained with L21-MTL. Both results were significantly (MEG: *p* = 0.000, EEG: *p* = 0.007) above chance level. For MEG, also pooling and L1-MTL yielded good results (66.5% and 67.8%, respectively) that were above chance level (pooling: *p* = 0.000, L1-MTL: *p* = 0.000). For EEG, L1-MTL yielded an average accuracy significantly better than chance (*p* = 0.01). Also, for both modalities within-subject decoding with MI training resulted in average accuracy better than chance (*p* = 0.000 for both MEG and EEG).

**Table 2.**
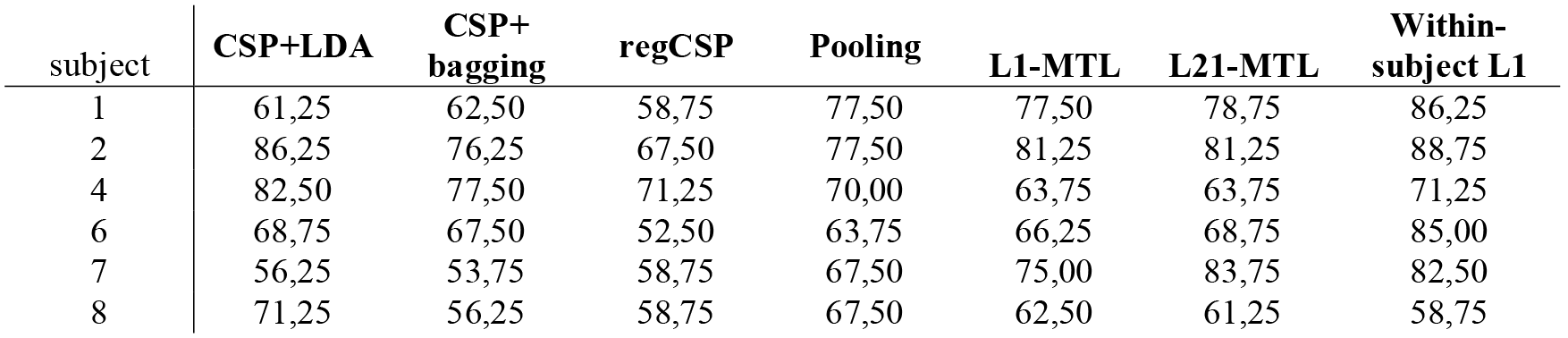
Accuracies of left-vs. right-hand MI decoding for the different methods using MI-trained MEG data.

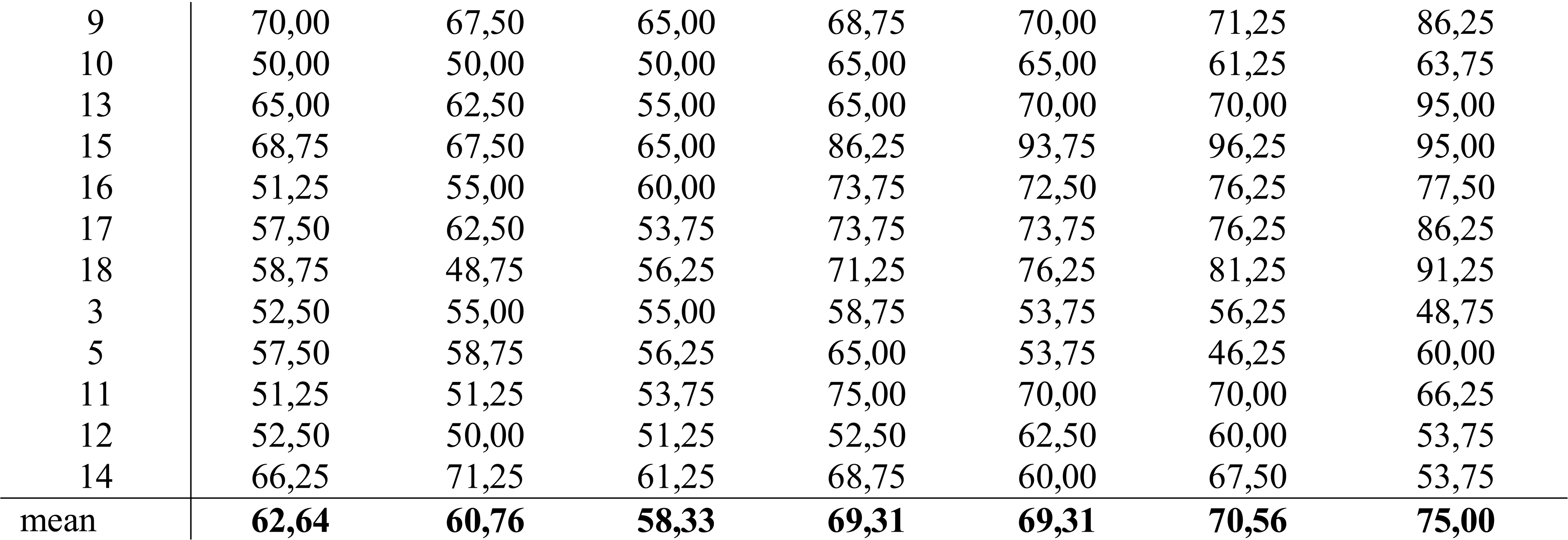

**Table 3.**
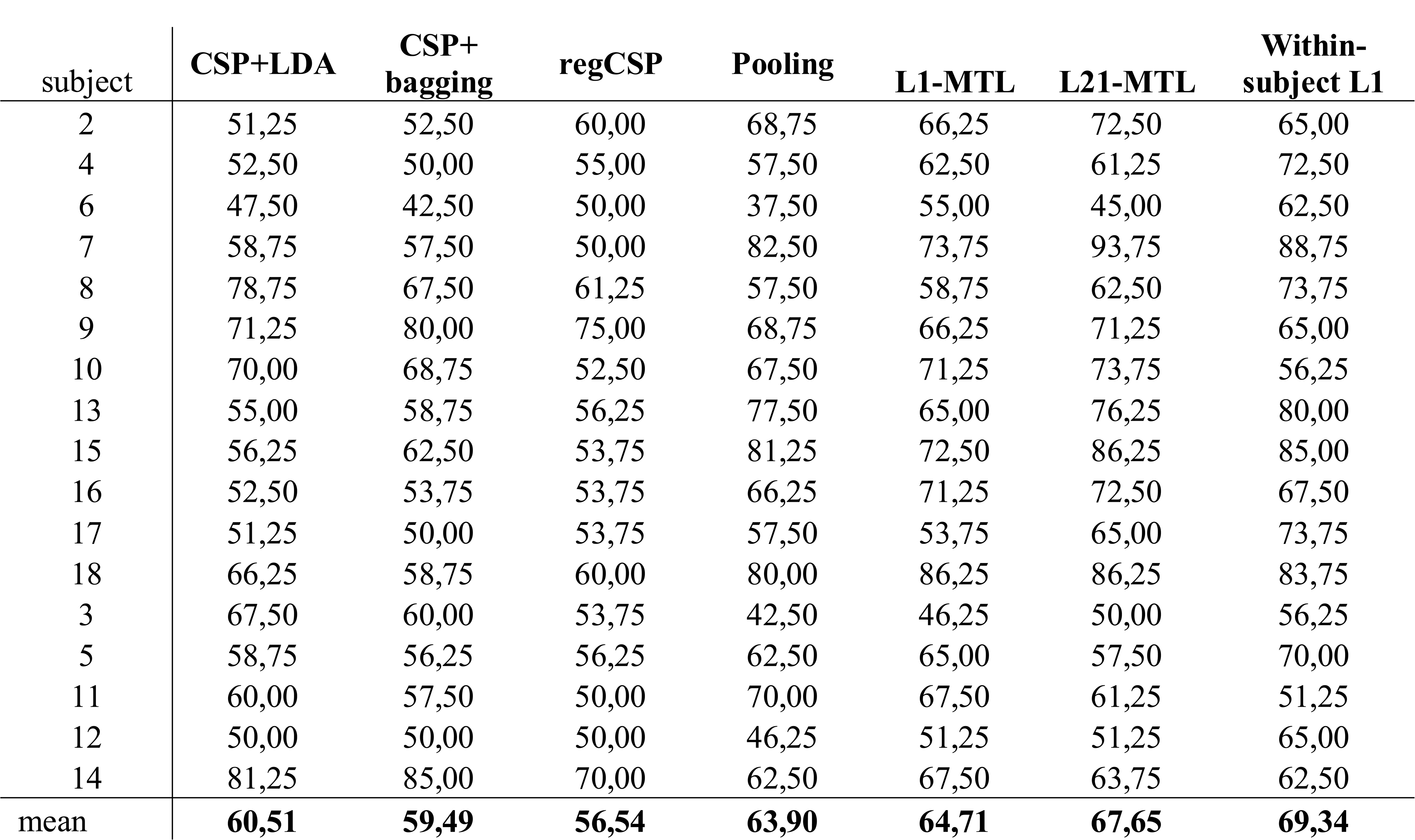
Accuracies of left-vs. right-hand MI decoding for the different methods using MI-trained EEG data.

The statistical differences between the methods (*p*-values, calculated by paired t-tests and Bonferroni-corrected for multiple comparisons) are presented in Supplementary Table S1 for MEG and in Supplementary Table S2 for EEG.

For MEG, Friedman’s test indicated significant effect of method on classification results (χ^2^ = 34.29, *p* = 0.000). The post-hoc multiple comparison test revealed that pooling, L1-MTL and L21-MTL were significantly better (*p* < 0.05) than the CSP-based methods. Also, no significant difference was found between within-subject L1 and inter-subject and L21-MTL or pooling. For EEG, Friedman’s test also indicated a significant effect of method on results (χ^2^ = 19.84, *p* = 0.003). In post-hoc multiple comparisons, it was found that L21-MTL was significantly better than CSP+bagging, regCSP and pooling. In this case, no significant difference was detected between within-subject L1 and L21-MTL, L1-MTL or pooling.

### 2.3. Offline training by passive movements

The classification results for MEG data with training by passive movements are presented in Table 4 and for EEG data in Table 5. For all subjects, the online within-subject MEG decoding yielded an average accuracy of 62.8%, which was not significantly better than chance. For EEG, the offline within-subject decoding resulted in an average accuracy of 66.8%, which was significantly above chance level (*p* = 0.008). For both MEG and EEG data, none of the inter-subject methods yielded accuracies significantly above chance level.

**Table 4.**
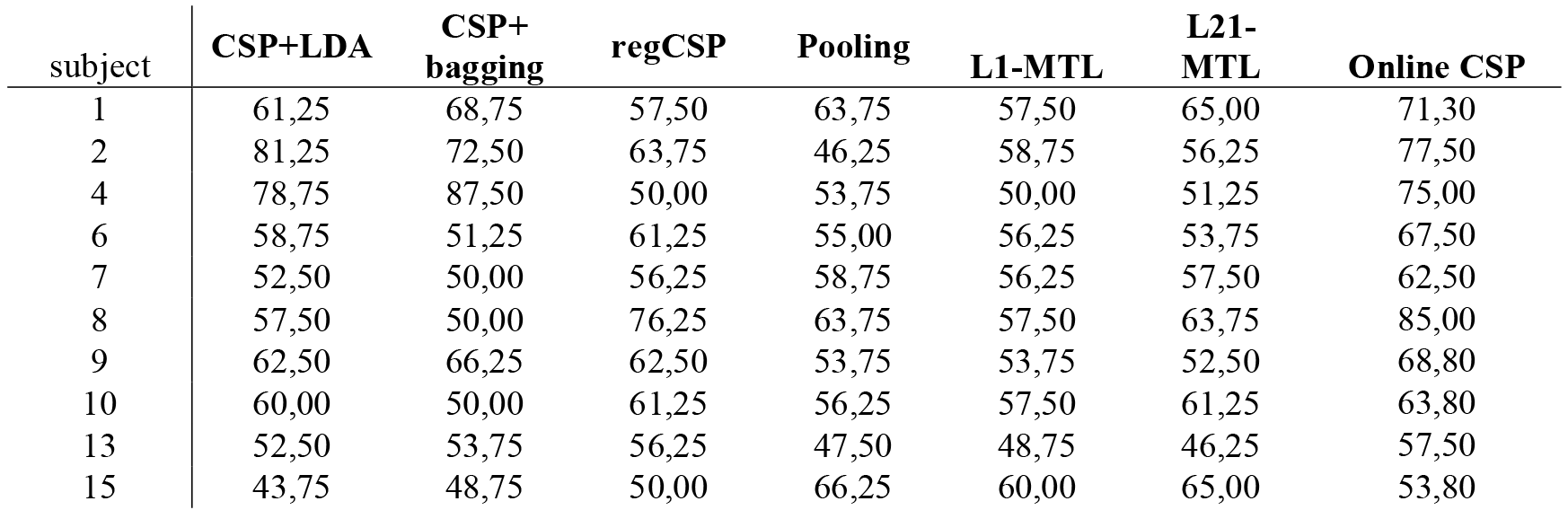
Accuracies of left-vs. right-hand MI decoding for the different methods using PM-trained MEG data.

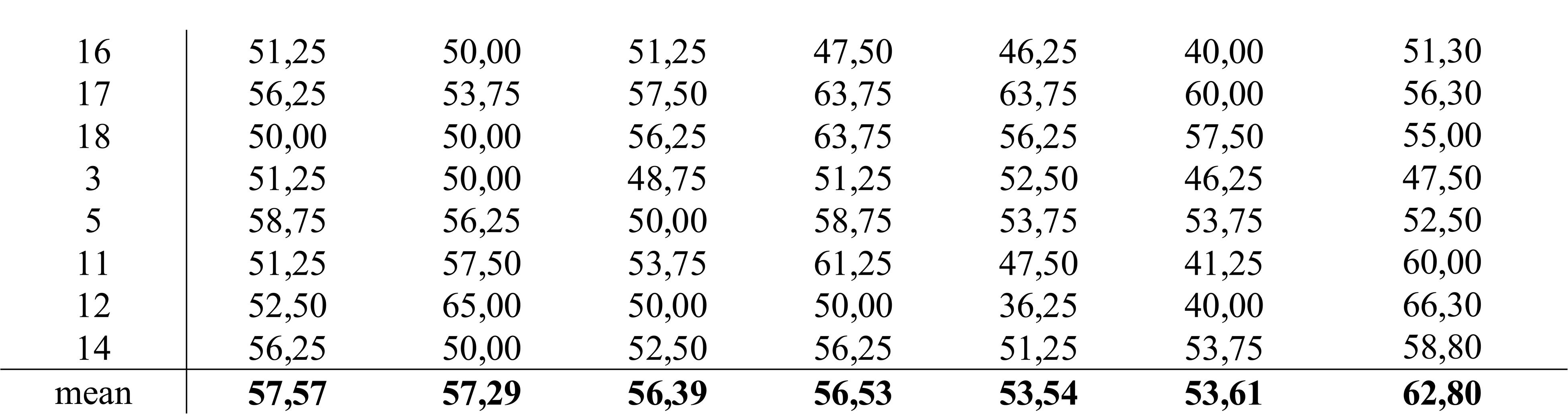

**Table 5.**
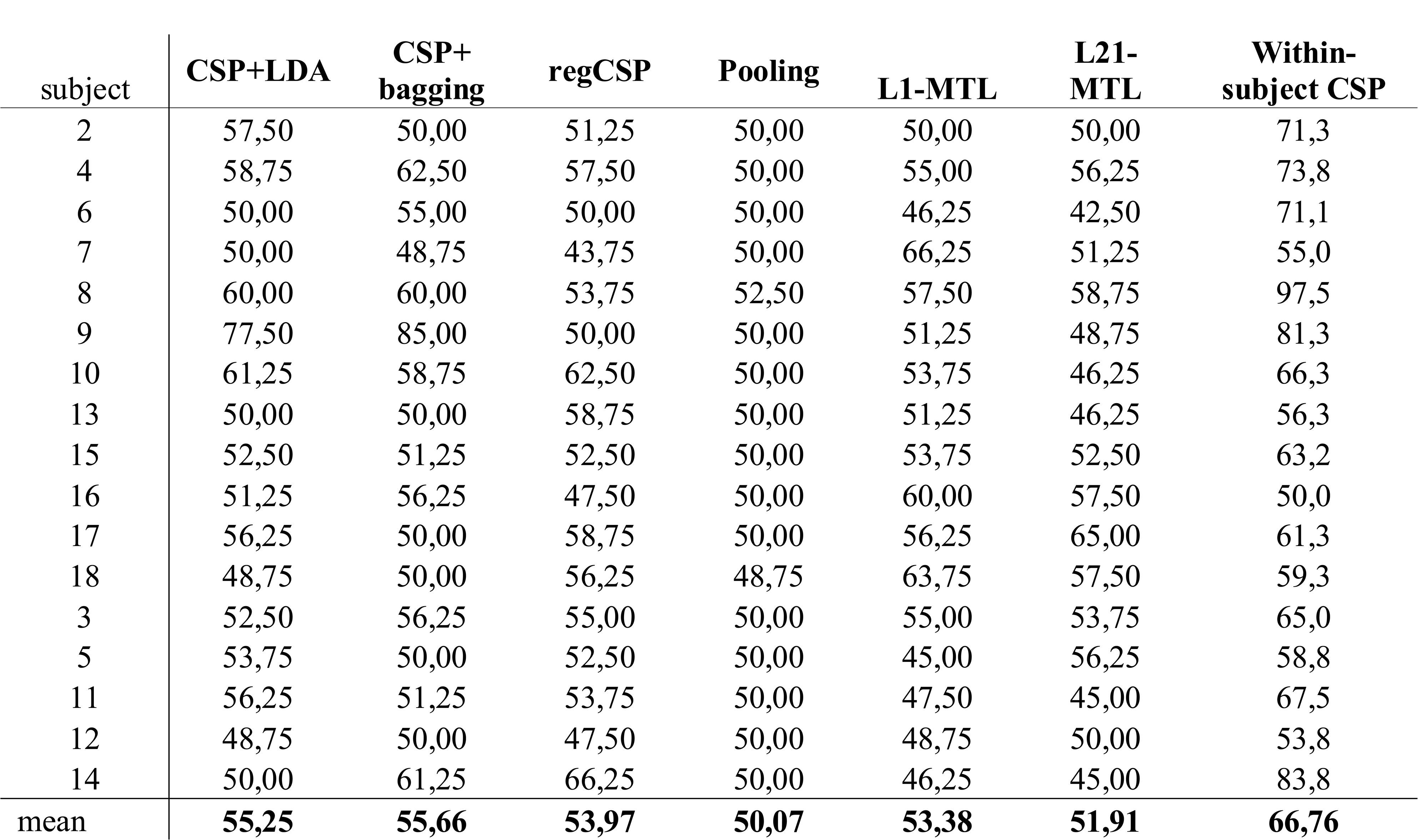
Accuracies of left-vs. right-hand MI decoding for the different methods using PM-trained EEG data.

The statistical differences between the methods (*p*-values, calculated by paired t-test, Bonferroni-corrected for multiple comparisons) are presented in Supplementary Table S3 for MEG and in Supplementary Table S4 for EEG.

For MEG, Friedman’s test showed a significant difference between methods (χ^2^ = 16.81, *p* = 0.01). The post-hoc multiple comparison test showed that within-subject decoding was significantly better than all inter-subject methods. For EEG, the results were quite similar; Friedman’s test indicated significant effect of method (χ^2^ = 34.21, *p* = 0.000) and the post-hoc multiple comparison test revealed that within-subject decoding was superior to all inter-subject methods. Also for EEG, pooling yielded results inferior to all other inter-subject methods.

## 3. DISCUSSION

In the current study, we investigated methods for inter-subject classification of motor imagery from MEG and EEG signals. We found that multi-task joint feature learning with logistic regression and l_2,1_-norm regularization is an effective method for decoding left- and right-hand MI from both MEG and EEG data. The accuracy of L21-MTL was better than that of the other inter-subject decoding methods both for MEG and EEG data, when other subjects’ MI was used for training. In addition, with this method we achieved accuracies significantly above chance level for both modalities. Also L1-MTL resulted in good average accuracy across subjects. When trained by MI, none of the CSP+LDA– based classification methods tested in this study reached an average accuracy equal to or higher than that of L21-MTL, L1-MTL or pooling with either MEG or EEG. Importantly, for both MEG and EEG, there was no statistical difference between within-subject L1 decoding and inter-subject L21-MTL decoding, suggesting that on average the performance of those methods was similar.

Besides the average accuracy across subjects, it is also important to consider how inter-subject decoding affected the performance of individual subjects. Many subjects showed poor performance during the online BCI task, when the decoder was trained using their own PM data. By using the subject’s own MI data and cross-validation, the accuracy was improved for most of the poor online performers. This offline analysis implies that the activity patterns of left- and right-hand MI were indeed discriminable in most subjects with low accuracy in the online task. It seems that PM training is not robust enough for MI classification, since there was large inter-subject variability in online decoding.

However, it is also noteworthy that some subjects have poor decoding performance because of insufficient SMR modulation. For example, Subjects 3 and 5 did not show any discernible SMR modulation during MI, and their accuracy remained under chance level even when their own MI was used for training. In case of the mentioned subjects, inter-subject decoding did not remarkably improve the results. On the contrary, for Subjects 11, 12 and 14, L21-MTL yielded better results with MEG (also with EEG for Subjects 11 and 14) compared to within-subject L1. We may thus conclude that for poorly-performing subjects showing at least some SMR modulation, inter-subject learning might improve the performance. On the contrary, for the good performers using other subjects’ training data were not beneficial, as most of them got worse results compared to training with their own data. However, inter-subject training still has the advantage of saving time in BCI sessions. It is an interesting question for future studies whether the good performers can adapt their brain activity such that they eventually get equally good accuracy with the generalized decoders.

As expected, training with PM resulted in lower inter-subject accuracy compared to training with MI – in fact, in this case the decoding accuracy was below chance level with all methods and both MEG and EEG. It is likely that this result is due to overfitting. It seems that the relevant features in PM data are different than the features in MI data, and thus it is difficult to classify MI using a classifier trained with other subjects’ PM. Some subjects achieved good performance when their own PM data were used for training, but the decoding accuracy decreased substantially when the training data were collected from other subjects. As a conclusion: even if the subject’s own PM pattern would be similar enough to the MI pattern to enable decoding of MI, the inter-subject generalized PM features are likely too different from the individual MI features. According to an EEG study by Formaggio an co-workers^31^, the activation patterns caused by MI are more lateralized than those caused by PM, which might explain this difference. However, the exact neural mechanisms underlying the difference between PM and MI remains unclear and should be investigated in the future. With regard to these findings, it can be argued that in inter-subject learning the performed mental task should be as similar as possible for the train and test subjects.

In their MEG classification study, Kia and co-workers ^17^ used a bootstrapping approach for dividing the data into different tasks, i.e. the samples for train and test datasets were randomly selected from a pool of data from different subjects. As samples for every task were collected from the same database, it is inevitable that MEG data samples from one subject are present in multiple tasks as well as in the test data. Therefore, the performance of the classifier may not reflect the true inter-subject generalizability. In the current study, we modified the approach of Kia and co-workers ^17^ and divided the data in discrete training and test datasets to ensure that no training data were collected from the test subject. It is noteworthy that the modified classifier still yielded sufficient inter-subject accuracy in MI classification. Kia and co-workers ^17^ used the L21-MTL method for classifying simulated MEG data and real MEG responses to visual stimuli, but to our knowledge the current study is the first one in which it was used to decode motor imagery.

According to our results, MTL seems to yield slightly but not significantly better results for MEG than EEG. In EEG we used fewer measurement channels and thus less features for classification than in MEG, which might explain the results. However, selecting an equal number of EEG electrodes and MEG sensors would not be neurophysiologically justified, as they would then cover a different area of the scalp. Therefore, we chose to select channels above the motor cortex in both modalities. It is also possible that MEG has a better signal-to-noise ratio than EEG, or that MEG is able to detect signals from sources not clearly visible to EEG. In light of these results, probably the best approach would be to utilize information from both MEG and EEG.

For the offline analyses, we wanted to have optimal data quality. Therefore, we suppressed interference in MEG with the signal-space separation method, aligned the head positions, and manually removed artifacts from EEG. It should be pointed out that this kind of thorough preprocessing is often not possible, or at least not practical, during online analysis. Head position alignment could be implemented in an online MEG data processing pipeline in the near future, so its use in the offline analysis was considered reasonable. In the case of EEG, one should carefully avoid artifacts during online decoding.

The motivation of decoding left-vs-right hand MI is based on the assumption that neural plasticity during stroke recovery might result in decreased functionality if inappropriate activation patterns are enhanced. Many stroke patients with motor disabilities have difficulties in SMR modulation in the affected hemisphere. Instead, the unaffected hemisphere shows increased motor-related activity due to reduced intracortical inhibition. Therefore, discriminating between MI and resting state is not likely a good target for a rehabilitative BCI, since it might only boost the activity of the motor areas on the healthy hemisphere instead of the affected hemisphere. Here we have shown that inter-subject decoding of left-vs-right MI from MEG and EEG yields reasonably good accuracy. The presented L21-MTL method can also be applied to online decoding.

In future studies, we aim at designing an adaptive online BCI paradigm based on the results of the current study. The BCI training could be started with generalized classifiers, and in the following sessions, as the patients’ performance improves, their own data could be added to the training set in order to update the classifiers, as has been proposed in recent studies ^32,33^. Another valid option is to use only the generalized classifier trained with other subjects. In this case, the test subjects should adapt their neural activity to match the MI representation of the healthy subject population. The latter approach would encourage the patients to change the activity of the damaged motor cortex towards normal function, which could boost rehabilitation. As the key concept in BCI therapy is to enhance activity-dependent brain plasticity^34,35^, it is crucial to ensure that the targeted activity corresponds to the normal motor-cortex function. Therefore, calibrating the BCI with healthy subjects’ data might actually lead to improved clinical outcome compared to calibration with patient’s own data. This issue will also be addressed in future studies.

## 4. METHODS

### 4.1. Participants

Eighteen healthy volunteers (5 males, 13 females, age 27.7 ± 5.0 y, 3 left handed by self-report) were recruited for this study. None of the subjects reported any neurological illnesses or motor deficits, and all subjects had normal or corrected-to-normal vision. Subjects having any metal objects inside their body or other MEG contraindications were excluded from the study. The study was approved by the Aalto University Research Ethics Committee. The research was carried out in accordance with the guidelines of the Declaration of Helsinki, and the subjects gave written informed consent prior to the measurements. Two of the subjects had participated in our previous experiment ^11^, and the others had no previous BCI experience.

### 4.2. Pneumatic hand stimulators

The stimulators used in this study consisted of two plastic frames supporting each hand, and of 8 elastic pneumatic artificial muscles (PAM; DMSP-10-100 AM-CM, diameter 10 mm, length of the contracting part 100 mm; Festo AG & Co, Esslingen, Germany) attached to fingers 2–5 of both hands with tape. During the experiment, the subject’s hands and arms were resting on the plastic frames and the PAMs were touching the fingertips horizontally from beneath the frame (Fig. 2). The PAMs were connected to the frame via moving hinges, so that the stimulator system could be adjusted to fit the subject’s hand.

**Figure 2.**
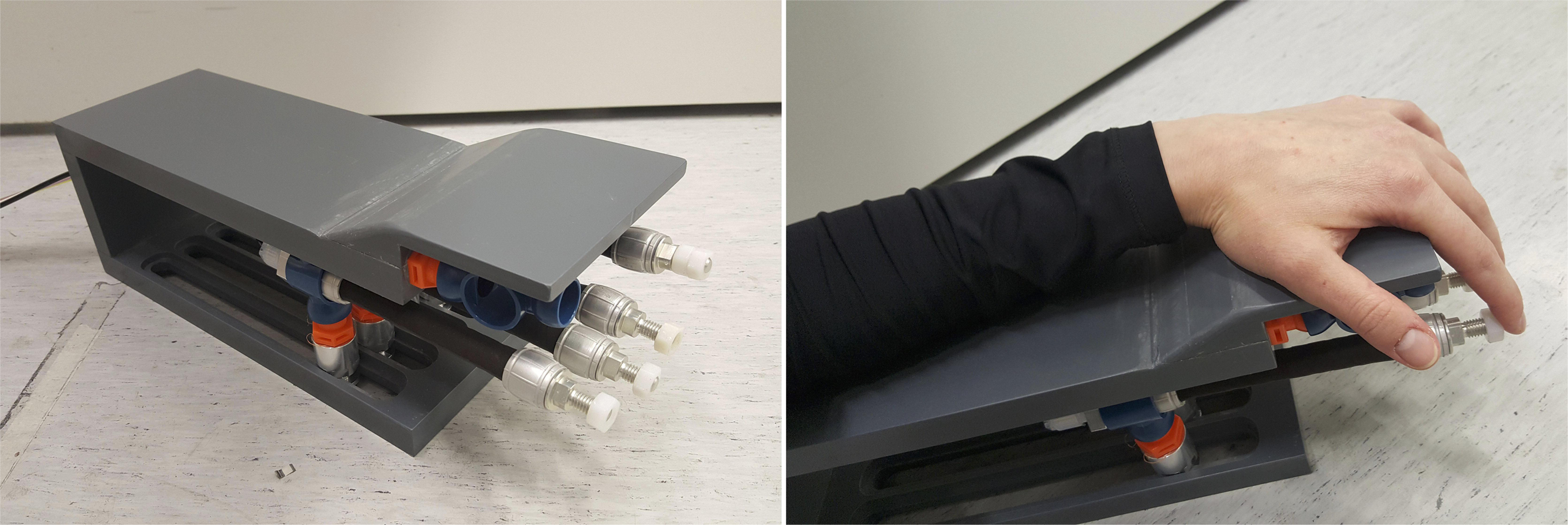
Pneumatic passive-movement stimulator.

During the stimulation paradigm, the PAMs moved horizontally according to the internal air pressure (1–4 bar). The pressure to each PAM was switched on and off by a solenoid valve (SY5220-6LOU-01F-Q, SMC Corporation, Tokyo, Japan) controlled by trigger pulses. The valves were placed outside the magnetically shielded room, and 3.5-m long semi-elastic tubes (internal diameter 2.5 mm) conveyed the pressurized air to the PAMs. A computer controlled the valves such that all the four PAMs for either left or right hand contracted sequentially (each for 500 ms) at intervals of 500 ms and flexed the fingers, thus mimicking small finger-tapping movement. After the contraction, each PAM returned to its resting length and extended the corresponding finger, returning it back to the initial positions. This four-finger movement sequence was repeated twice during each epoch.

### 4.3. MEG and EEG recordings

MEG was recorded at the MEG Core of Aalto University School of Science with a 306-channel Elekta Neuromag™ Vectorview system (Elekta Oy, Helsinki, Finland) using a sampling frequency of 1 kHz. Five head-position indicator (HPI) coils were attached to the subject’s scalp for head position estimation and alignment with a standard coordinate system. Visual stimuli were delivered on a screen located approximately 50 cm from the subject’s eyes via a projector outside the shielded room. EEG was measured simultaneously using a MEG-compatible cap with 60 electrodes and 1-kHz sampling rate. However, only the MEG data were used for real-time classification.

Raw MEG data were continuously written in 300-ms segments to a network-transparent ring buffer ^36,37^ hosted by the MEG acquisition workstation (6-core Intel Xeon CPU at 2.4 GHz, 64-bit CentOS Linux, version 5.3). The buffer was read over a local network connection by another computer (Intel Core i7-4771 CPU at 3.5 GHz, 64-bit Ubuntu Linux, version 14.04). This computer processed the MEG data in real time using functions implemented in the MNE-Python software ^38^, controlled the pneumatic hand stimulators and presented the visual stimuli using PsychoPy version 1.83 ^39^.

### 4.4. Experimental paradigm

Each measurement session consisted of the two tasks described below. The whole experimental paradigm is illustrated in Fig. 3. The subjects were allowed to rest eyes closed between the tasks for a few minutes. The duration of each task was approximately 15 min, and the whole experiment lasted for approximately 40 min including breaks.

**Figure 3.**
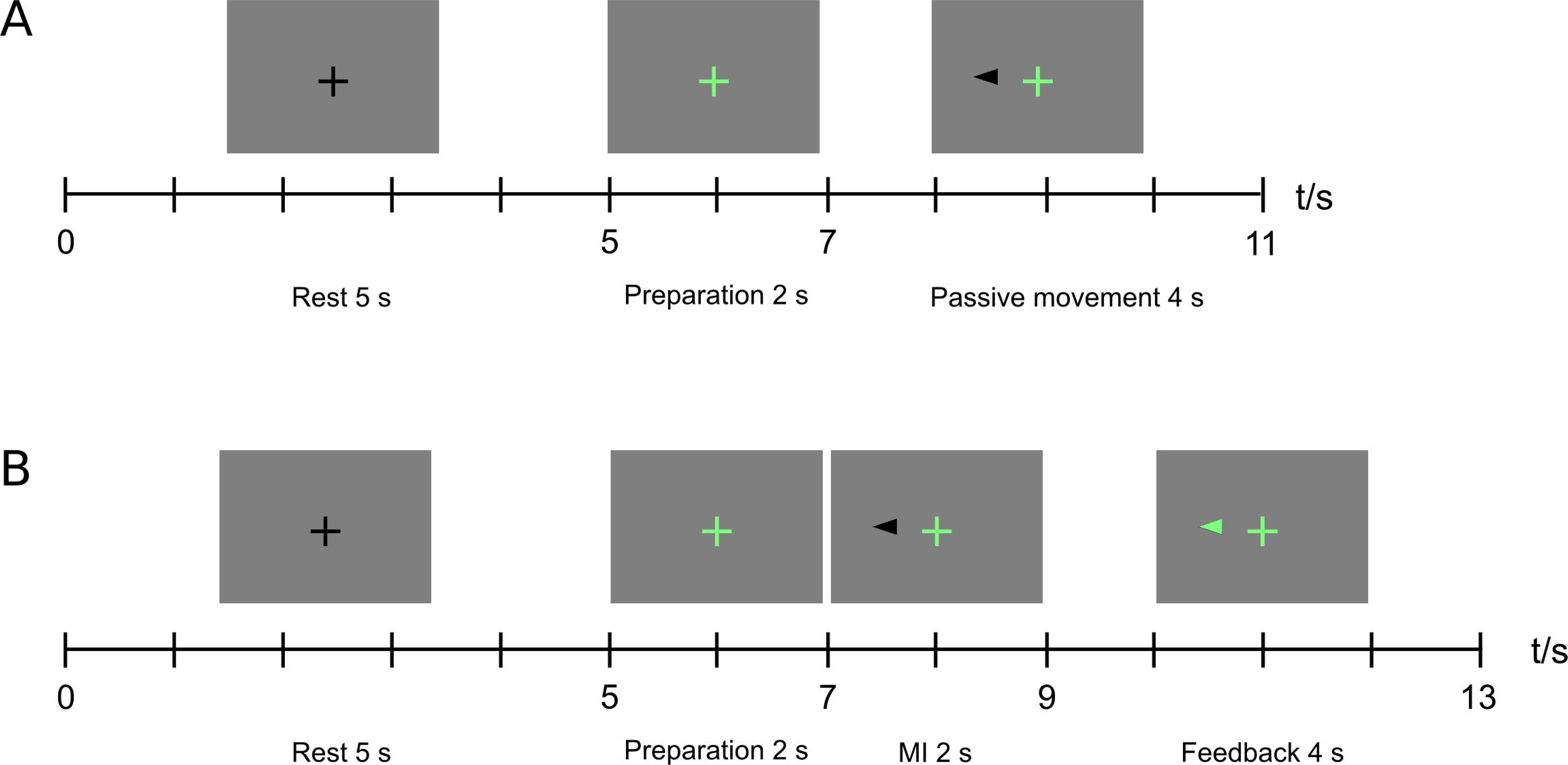
The experimental paradigm in (A) passive movement, (B) motor imagery tasks.

In the passive-movement task, hand movements were elicited by the pneumatic stimulators according to the predefined sequence. Each sequence onset was accompanied with a visual cue. The subjects were instructed to keep their muscles relaxed and gaze on a fixation cross throughout the task. A total of 80 passive-movement epochs, 40 for each hand in a pseudo-randomized order, were delivered. Each epoch began with a gray background and a black fixation cross in the middle of the screen. At second 5, the fixation cross turned green, indicating preparation for the upcoming movements. At second 7, a black cue arrow appeared on either left or right side of the screen, and the stimulators moved the subject’s fingers on the corresponding hand continuously for 4 s. At second 11, a black fixation cross was shown and the PAMs returned to their original positions, leaving the fingers at rest.

After the first task, the subject was instructed to relax for a few minutes without making large head movements. In the meantime, a classifier was trained using the MEG data collected during passive movements. Signals from 64 MEG gradiometers over the parietal areas were included in the online analyses (see Fig. 4). First, the frequency band showing the strongest SMR suppression was estimated by comparing the SMR power at rest, i.e. during the pauses in the first experiment, and during passive movements. Power decrease was calculated in 1-Hz frequency bins in 8–30-Hz range. The center frequency was defined as the frequency with the highest power decrease, and the frequency band of interest as the center frequency ± 3 Hz. After that, relevant signal features were extracted with SSD^24^ and consecutive Common Spatial Patterns (CSP) filters. The frequency band of interest was used for SSD. Other details for SSD and CSP can be found in our previous study^11^. Finally, a Linear Discriminant Analysis (LDA) classifier was fit to the data to discriminate between left- and right-hand-imagery-related suppression of the SMR.

**Figure 4.**
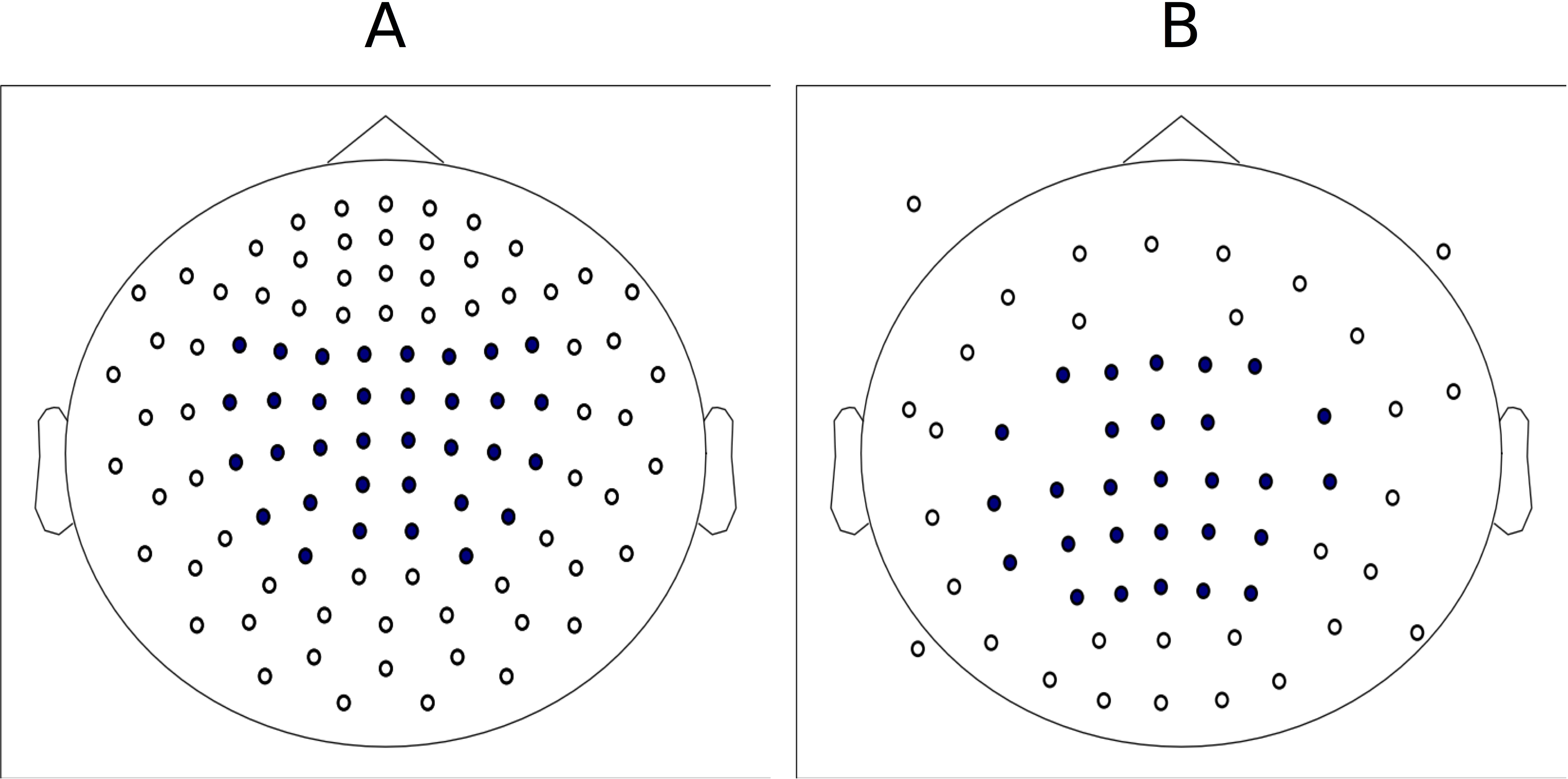
Locations of (A) MEG sensors and (B) EEG electrodes (marked in blue) included in analyses.

In the motor imagery (MI) task, the subjects were instructed to imagine finger movements similar to the passive movements they received in the earlier task and to pay attention to the visual and tactile feedback following each MI period. The aforementioned SSD and CSP filters and the LDA classifier were applied in real time to each MI epoch to predict left- vs. right-hand MI.

The MI task comprised 80 trials of MI with feedback. Each trial began with a black fixation cross in the middle of the screen. At second 5, the fixation cross turned green, indicating preparation for the upcoming movement. At second 7, a black cue arrow appeared on either side of the screen, indicating the beginning of MI. At second 9, the feedback started; if the MI trial was successful, i.e. the classification probability of the target hand exceeded 50%, the cue arrow turned green and the predefined sequence of PAM stimulator movements started, lasting 4 s. In case of an unsuccessful MI trial, the cue arrow turned red and the PAMs did not move.

### 4.5. Preprocessing for offline analyses

Similar preprocessing routines were applied to the data from both tasks. First, the data were processed with the MaxFilter software (version 2.2.15; Elekta Oy, Helsinki, Finland)^40^, including transformation of head positions of all subjects into the same orientation, signal space separation for suppressing external magnetic interference and removal of static bad channels. The common-average reference was utilized for all analyses of EEG data. Both MEG and EEG data were bandpass-filtered to 6–45 Hz with a fast Fourier transform (FFT)- based finite impulse response (FIR) filter, with filter length equal to the epoch length. The data were split into 3-s epochs and baseline corrected (epoch −1.0…2.0 s, baseline −1.0…0.0 s from cue onset). Artifact correction for EEG data was done by performing independent component analysis (ICA; implemented in MNE-Python^38^) and manually removing components including artifacts or high-amplitude noise. 64 MEG sensors and 28 EEG electrodes close to parietal areas were included in further analyses (see Fig. 4).

### 4.6. Train and test data in offline analyses

The measurements were split into train and test data in a leave-one-subject-out manner, i.e. the data from all except one subject were used for training the classifiers and the remaining subject’s data for testing classification performance. The MEG and EEG signals were evaluated separately. In addition, we evaluated the validity of both passive movements (PM) and MI of other subjects for training the classifier. In case of PM, the training data were PM epochs from all but one subject and the test data were MI epochs from the remaining subject.

### 4.7. Feature extraction and classification

#### 4.7.1. Spatial filtering and LDA

Before CSP filtering, the data dimensionality was reduced with spatio–spectral decomposition (SSD)^24^. The frequency band of interest was defined as 10 ± 3 Hz in both datasets for all subjects, and 20 SSD components were retained. CSP decomposition was calculated on the SSD-filtered data. Ten CSP components were extracted, averaged over time and log-transformed, and finally used as features for the LDA classifier.

In the CSP+bagging approach, the LDA classifier was first fit to each training subject’s data separately, and the individual predictions were aggregated to form the final predictions for the test data. The BaggingClassifier algorithm of the scikit-learn Python package ^41^ was modified such that instead of dividing the training data into random subsets, each division contained all data from one train subject.

The regularized CSP algorithm was adopted from Regularized CSP (RCSP) Matlab toolbox ^21^ and implemented in Python. The generic covariance matrices **G**_*c*_ for each class *c* were calculated as weighted sums of covariance matrices **C**_*c*_ from *N* training subjects ^22^:

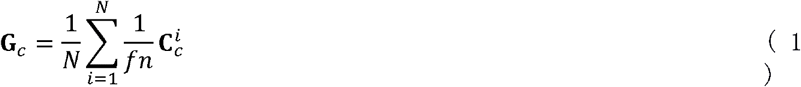

where *f* is the Frobenius norm:

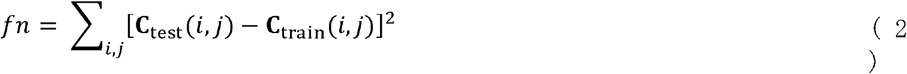

The objective function of RCSP was formulated as [Lotte & Guan 2010]:

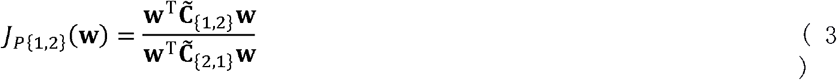

where 
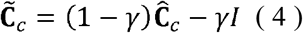
and 
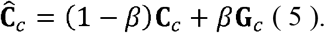

Values for parameters *β* and *γ* were selected by cross-validation such that one subject was used for testing, half of the remaining subjects for parameter optimization and the other half for training. The parameters were optimized discretely on range from 0 to 1 with a step size of 0.1. Leave-one-out cross-validation was used for the parameter optimization dataset to find out the parameters yielding the best classification results, and those parameters were used for the actual training and testing.

### 4.8.2. Multi-task joint feature learning

The MTL optimization problem can be expressed as an empirical risk minimization problem:

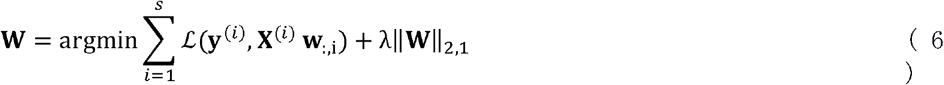

where ║**W**║ is the l_2,1_ norm for matrix **W**:

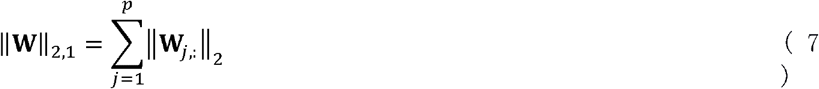

The multi-task logistic regression minimization problem can thus be formulated as:

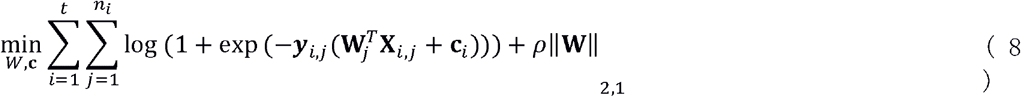

where **X**_*i,j*_ denotes sample *j* of the *i*-th task, **y**_*i,j*_ denotes the corresponding label, **W**_*i*_ and **c**_*i*_ are the model for task *i* and the regularization parameter *ρ*_1_ controls group sparsity.

Prior to feature selection and classification, the epochs were decimated by 10, i.e. every 10^th^ timepoint was selected, to speed up calculation. The bootstrap data sampling scheme used in the L21-MTL algorithm of MTJFL toolbox was replaced by a leave-one-subject-out scheme, in which no data from the test subject were included in the training set. In the MTL, the data were split such that each task included all data from a single train subject, and the test data included all data from the test subject. In the pooling approach, the data from all except one subject were used for training, and the data from the remaining subject for testing. This train/test split ensured that the classifier actually generalized between the subjects and did not use any of the test subject’s data for training. In the within-subject decoding, 10-fold cross-validation was used for dividing each subject’s data into train and test sets. In all approaches, the regularization constant lambda was selected by choosing the parameter yielding the best results in L21-MTL classification, and a similar lambda was used also for L1-MTL. The parameter was selected separately for all datasets and conditions.

### 4.9. Statistical analysis of classification results

The chance level for left-vs-right MI classification was defined according to the study by Combrisson and Jerbi^30^. The probability for getting a correct classification by chance is 58.7% for 80 samples and a *p*-value of 0.05, and we used this value as a threshold for determining whether the decoding results differ from chance level.

The statistical difference between results of different classification approaches was calculated with Friedman’s test, a nonparametric version of two-way analysis of variance. Post-hoc analysis was conducted using Bonferroni correction for multiple comparisons, implemented in MATLAB (R2017a; Mathworks Inc., MA, USA) function *multcompare*.

### 4.10. Data availability

The data measured and analysed for the current study are available from the corresponding author on request.

## Acknowledgements

The study was supported by Emil Aaltonen foundation (H.-L.H.) and Academy of Finland grant #295075 “Neurofeed” (L.P.).

## Author contributions

H.-L.H and L.P. designed the studies. Data were collected by H.-L.H. Data analyses were carried out by H.-L.H. The manuscript was written by H.-L.H. and L.P. Figures were prepared by H.-L.H. Both authors reviewed and accepted the manuscript.

## Additional information

The authors declare no competing interests.

## Supplementary Information

See the attached file.

